# *geck*: trio-based comparative benchmarking of variant calls

**DOI:** 10.1101/208116

**Authors:** Péter Kómár, Deniz Kural

## Abstract

**Motivation:** Classical methods of comparing the accuracies of variant calling pipelines are based on truth sets of variants whose genotypes are previously determined with high confidence. An alternative way of performing benchmarking is based on Mendelian constraints between related individuals. Statistical analysis of Mendelian violations can provide truth set-independent benchmarking information, and enable benchmarking less-studied variants and diverse populations.

**Results:** We introduce a statistical mixture model for comparing two variant calling pipelines from genotype data they produce after running on individual members of a trio. We determine the accuracy of our model by comparing the precision and recall of GATK Unified Genotyper and Haplotype Caller on the high-confidence SNPs of the NIST Ashkenazim trio and the two independent Platinum Genome trios. We show that our method is able to estimate *differential* precision and recall between the two pipelines with 10^-3^ uncertainty.

**Availability:** The Python library *geck*, and usage examples are available at the following URL: https://github.com/sbg/geck

**Supplementary information:** Supplementary materials are available at *bioRxiv*.

## 1 Introduction

Next-generation sequencing (NGS) has enabled entire human genomes to be sequenced in a single experiment and has, since its development, played a crucial role in many areas including population genetics (Auton *et al.*, 2015; Mallick *et al.*, 2016), disease genetics (Bamshad *et al.*, 2011), and the study of human mutations (Veltman and Brunner, 2012). Many variant callers have been developed to call genetic variants from NGS data (Sandmann *et al.*, 2017), employing multiple steps of data processing and statistical inference (Li, 2011). Due to their wide impact, benchmarking variant calling pipelines is of utmost importance (Li, 2014; Shringarpure *et al.*, 2015).

A widely-used method of measuring the accuracy of variant calling pipelines is tallying mis-called variants on samples that have been characterized previously (Olson *et al.*, 2015; Hwang *et al.*, 2015; Cornish and Guda, 2015). High-confidence variant sets, or “truth sets”, are compiled by teams of experts, such as the Genome in a Bottle (GIAB) Consortium, by integrating variant calls from several sequencing and data processing technologies. Due to careful curation, these data sets reach very high fidelity, and provide a benchmarking reference for SNPs, indels (Zook *et al.*, 2014; Talwalkar *et al.*, 2014; Boutros *et al.*, 2014), and structural variants (Parikh *et al.*, 2016; Fang *et al.*, 2016; Nutsua *et al.*, 2015). In all cases, high-coverage sequencing data from a handful of (often related) individuals are required to produce a truth set (Zook *et al.*, 2016; Eberle *et al.*, 2017). Such truth sets are expensive to create, specific to the population, and biased toward the methods that produced them.

Sequencing data from related individuals provide an alternative way to detect errors made by a variant calling pipeline. Since de-novo mutation rates (~10^−8^ per locus (Veltman and Brunner, 2012)) are several orders of magnitude smaller than calling and genotyping error rates (often 10^−2^ – 10^−5^), one can use the variant calls inconsistent with the rules of Mendelian inheritance to assess genotyping errors. While not all errors result in a Mendelian violation, which limits the sensitivity of this strategy, as noted by Li (2014), additional assumptions about the error probabilities permit the estimation of genotyping error rates from parent-offspring dyads (Haaland and Skaug, 2013), siblings (Wang, 2004; Johnson and Haydon, 2007; Korostishevsky *et al.*, 2009), trios (Sobel *et al.*, 2002; Douglas *et al.*, 2002; Saunders *et al.*, 2007; Jostins, 2011) and larger pedigrees (Eberle *et al.*, 2017). If sequencing reads from many related individuals are available, recombination events can be inferred and used to estimate locus-specific genotyping errors (Markus *et al.*, 2011; Chen *et al.*, 2013; Peng *et al.*, 2013; Kojima *et al.*, 2013). All these approaches have the advantage of not relying on a truth set, thereby eliminating the bias towards established variant callers.

When modeling genotyping calling errors in related individuals, one is tempted to parametrize the genotyping error profile with a couple of parameters and estimate their values independently for different variant callers. This method has two significant shortcomings by neglecting the following correlations: First, errors made by different variant calling pipelines are often correlated due to identical assumptions (e.g. well-mapped reads) which, if violated in the input data, make mistakes of the pipelines coincide at particular loci. Second, mapping and calling errors depend on sequence context, which is similar among closely related individuals and induces correlations between genotyping errors in samples of different family members. To the extent of our knowledge, there exists no published method that addresses both correlations.

Here we present a statistical model that addresses both shortcomings for the case of a family trio. First, we aggregate the number of different genotype trios for the two pipelines jointly, and model this data. This way, we take correlations between two variant callers explicitly into account. Second, our model consists of a statistical mixture of different genotyping error profiles. Although each error profile assumes independence between family members, enabling an efficient mathematical solution, their mixture captures the correlations.

We validate our model in a series of experiments on real trios. First, we run two variant caller pipelines on publicly available NGS data from trios, and let our model estimate the precision and recall of the two pipelines. Then we compare these estimated performance metrics with their true values. The truth is directly calculated by comparing the calls on the children with their correct genotypes from the truth data sets. We demonstrate that our model is able to detect *differential* performance between two pipelines as small as ~ 10^−3^. With this level of accuracy, our model enables informed development of pipelines targeting less-studied variants and populations for which no curated truth set is available.

The rest of the paper is divided into three parts: In section 2, we introduce our notation, describe the structure and assumptions of our model, and show how one can use it to infer the number of genotyping errors and stratify them by type. In section 3, we describe how we validate our method on publicly available trio data. Finally, in section 4, we present the results of the validation experiment.

## 2 Model

### 2.1 Definitions and notation

**Trio genotypes.** Let *I* = {00,01,11} denote the set of individual, bi-allelic, unphased genotypes: homozygous reference (00), heterozygous (01) and homozygous alternate (11). Let *g* = {*g*_*p*_ ∈ *I* : *p* = 1, 2, 3} be an ordered trio of individual genotypes, representing the genotypes of variants in the genomes of father (*g*_1_), mother (*g*_2_) and child (*g*_3_). Let *T* denote the set of mathematically possible genotype trios, i.e. *T* = *I*^×3^, and let *t* ⊂ *T* denote its subset that conforms with Mendelian inheritance, i.e. *t* = {*g* ∈ *T* : (*g*_3,1_ ∈ *g*_1_ and *g*_3,2_ ∈ *g*_2_) or (*g*_3,1_ ∈ *g*_2_ and *g*_3,2_ ∈ *g*_1_)}, where *g*_3,1_ and *g*_3,2_ are the first and second symbols (0 or 1) of *g*_3_. One can show that |*T*| = 27 and |*t*| = 15, by enumeration.

**Observed genotypes.** Two variant calling pipelines run independently on samples from the three family members produce six separate sets of variant calls. We aggregate the counts of different genotype trio pairs of one class of variants (e.g. SNPs, short indels, structural variants) called by the two pipelines across all loci. This yields the joint counts of genotype trios 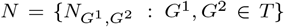, where 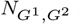 is the number of variants whose trio genotypes are called *G*^1^ by pipeline 1, and *G*^2^ by pipeline 2. An examples is shown in Figure 1.

**Fig. 1.**
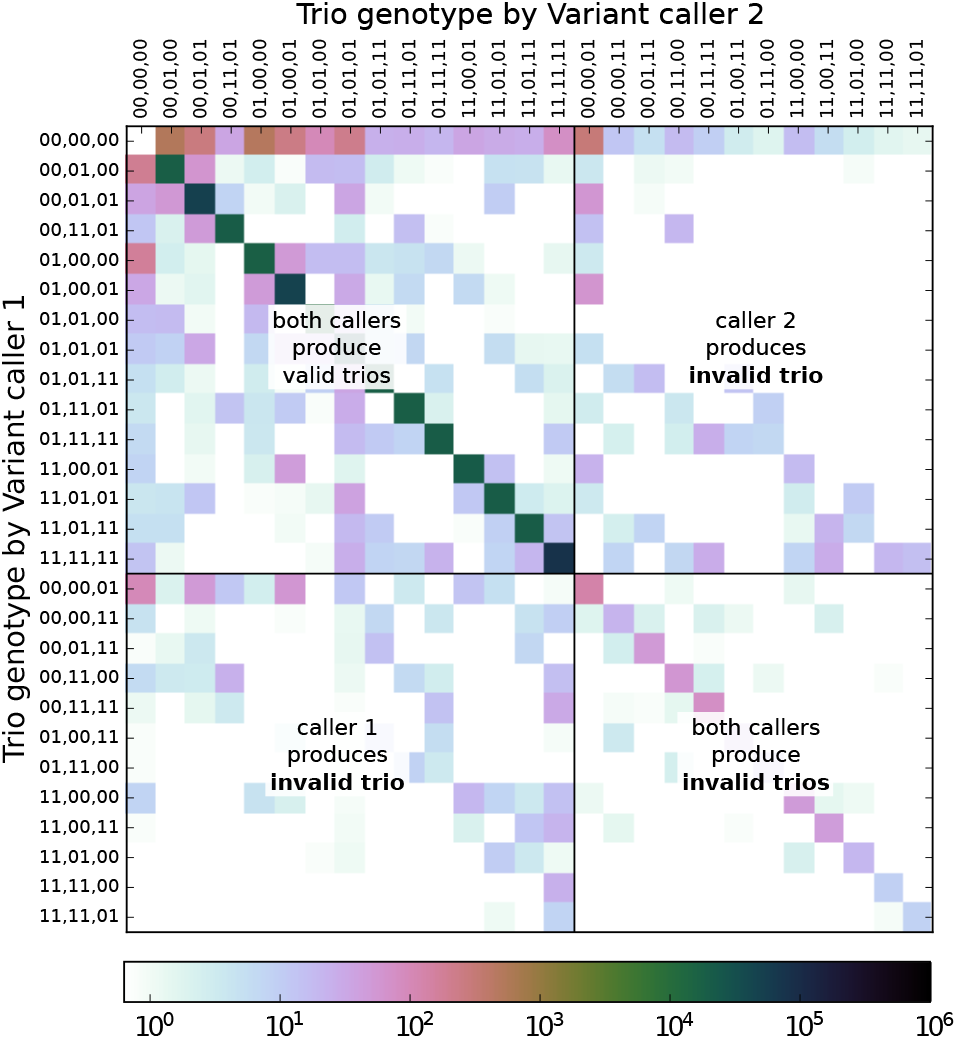
An example of the joint counts of observed trio genotypes *N*. Each cell at row *G*^1^ and column *G*^2^ counts the number of variants for which pipeline 1 calls *G*^1^ and pipeline 2 calls *G*^2^ trio genotypes. The 27 trio genotypes (*T*) are sorted such that the first 15 comply with Mendelian inheritance (*t*). Most variants populate the main diagonal, indicating strong correlation between the pipelines. If the pipelines could exploit that the three individuals form a trio, none of them would call invalid trios and only the upper left corner of the matrix would contain counts. Still some of the counts would be placed off the main diagonal, because the pipelines would not always agree.

**Correct genotypes**. Explicit information about the correct trio genotypes *g* ∈ *t* is missing from the observed counts *N*. Let 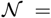 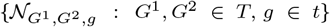 denote the (hidden) complete distribution of called and correct genotype trios, where 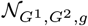 is the number of variants with called genotype trios *G*^1^, *G*^2^ and correct genotype trio *g*, such that 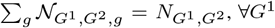. Inthe next section, we explicitly model the genotyping errors in order to infer the hidden components in 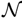, examples of which are shown on Supplementary Figures S.3, S.4, and S.5. Uncovering this data, 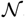, from the observed mixture, *N* is our main goal.

**Benchmarking metrics**. If obtained, the complete distribution 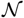 contains all necessary information to directly evaluate any benchmarking metric. We will use the notation 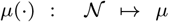 to denote any benchmarking metric, derivable from 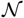. One important metric is the genotype confusion matrix, 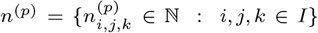, for individual *p*, where

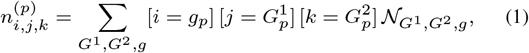

where [*x* = *y* is 1, if *x* = *y* and 0 otherwise. 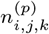 counts how many variants of person *p*, with correct genotype *i* ∈ *I*, are called *j* ∈ *I* by pipeline 1, and *k* ∈ *I* by pipeline 2.

Genotyping can be understood as a classification task with one negative (00) and two positive classes (01 and 11). For person *p*, we define the true positive, false negative, false positive and “mixed-up positive” counts for pipeline 1 as 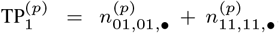,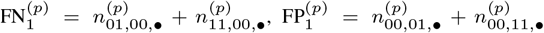 and 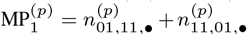, respectively, where • denotes summation over the third index. For pipeline 2, similar formulas apply where the summations are prescribed for the second index. We define precision and recall with the usual formulas: prec = TP/(TP + FP + MP) and rec = TP/(TP + FN + MP), for each person and each pipeline.

### 2.2 Generative model

We introduce a generative model in a form of a mixture of categorical distributions over (*G*^1^,*G*^2^) ∈ *T* × *T* pairs, where each mixture component is labeled by a correct trio genotype *g* ∈ *t*, and an error category *m* (defined in this section). The graphical representation of this model is shown in Figure 2. We define the parameters *f*, *θ*, *E* below.

**Fig. 2.**
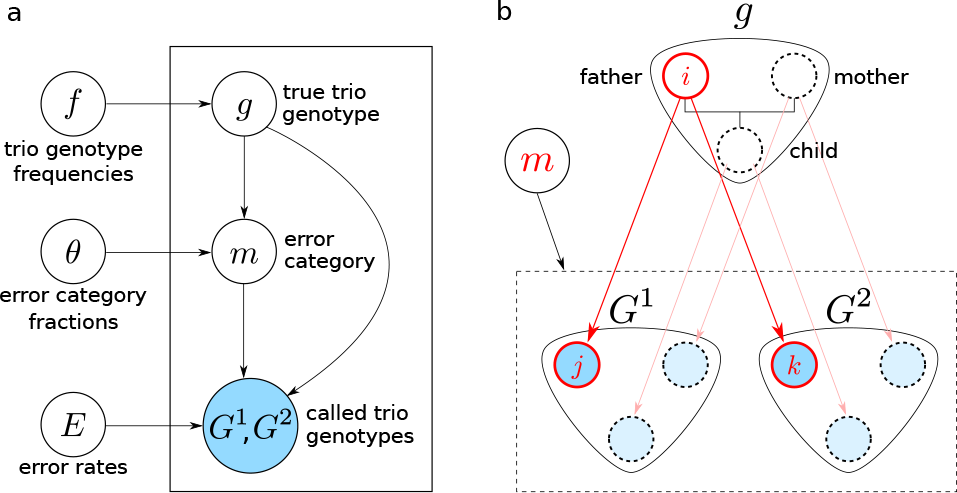
Graphical representation of the model. **a.** Parameters, *f*, *θ*, *E*, determine the distributions of hidden variables: correct genotype trio *g* and error category m, and the observed genotype trios: *G*^1^ and *G*^2^. **b.** Transitions from correct genotype trio *g* to observed genotype trios (*G*^1^;*G*^2^) are caused by genotyping errors. The correct genotype *i* ∈ *I* of a variant in the genome of each family member undergoes a transition, subject to the same error categorym, resulting in called genotypes (*j*, *k*) ∈ *I* × *I*.

**Correct frequencies.** Let *f* = {*f*_*g*_ : *g* ∈ *t*} denote the frequencies of each correct genotype trio *g* among the variants under investigation, i.e.

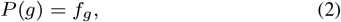

where *f* is subject to normalization, i.e. ∑_*g*_ *f*_*g*_ = 1. By treating all 15 components of *f* as independent parameters, our model is able to fit aggregate data of variants not showing Hardy-Weinberg equilibrium (a limitation of the models used by Haaland and Skaug (2013) and Douglas *et al.* (2002)). This way of parameterizing the correct frequencies makes our model more flexible than previous approaches that assume global allele frequencies (Haaland and Skaug, 2013;Wang, 2004; Johnson and Haydon, 2007; Browning and Browning, 2013; Douglas *et al.*, 2002; Saunders *et al.*, 2007; Korostishevsky *et al.*, 2009).

**Error categories.** We assume that each variant is affected by one of the error categories *M* = {*a*; *b*; *c*; *d*; *e*}: (*a*) Both pipelines are correct, (*b*) Pipeline 1 is correct, (*c*) Pipeline 2 is correct, (*d*) Pipelines agree (but not necessarily correct), and (*e*) Pipelines are arbitrarily dependent. Let *θ* = {*θ*_*g*,*m*_ : *g* ∈ *t*; *m* ∈ *M*} denote the fractions of variants, with correct genotype trio g, that are affected by error category *m*, i.e.

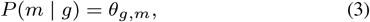

where *θ* is subject to normalization, i.e. ∑_*m*_ *θ*_*g*,*m*_ = 1; ∀*g*. The error category *m* is assumed to be the same for all three genomes at a given locus. Now, we extend the definition of 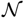 with this additional index, 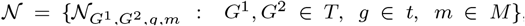, where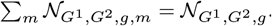. The idea of alternative error categories is used in Korostishevsky *et al.* (2009) and Heid *et al.* (2008), but none considered a mixture over error categories.

**Error rates.** Let 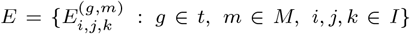 denote the probabilities of calling the genotypes (*j*,*k*), when the correct genotype is *i* for a variant whose correct genotype trio is *g* and is subject to error category *m*. Under a given *g* and *m*, we assume that errors happen independently, and with the same rate, for all members of the trio. In practice, this requires that the samples are prepared, sequenced and analyzed in the same way, separately from each other. This allows us to write the probability of each type of genotyping event 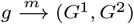 as the product

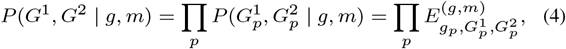

where *E* is subject to the normalization 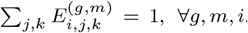. Different error categories *m* ∈ *M* impose different restrictions on the values of

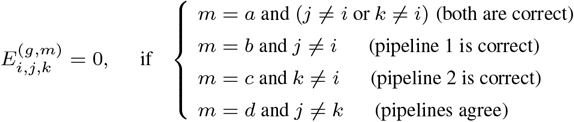

and *m* = *e* imposes no restrictions. Previously published models either assume an error rate independent of the correct genotype (Sobel *et al.*, 2002; Saunders *et al.*, 2007; Jostins, 2011; Browning and Browning, 2013) or model the *E* matrix with a few parameters (Wang, 2004; Johnson and Haydon, 2007; Korostishevsky *et al.*, 2009; Douglas *et al.*, 2002; Heid *et al.*, 2008). It is worth emphasizing that we only assume *conditional* independence of the errors in different family members (conditioned on fixed *g* and *m*). The marginal probability *P*(*G*^1^, *G*^2^) = ∑_*g*,*m*_ *P*(*G*^1^,*G*^2^,*g*,*m*) does not factorize under our assumptions, enabling our model to capture correlated errors across family members.

### 2.3 Limiting complexity

The model described so far has 480 algebraically independent parameters (See Supplementary A.1.1), which is comparable to 27 × 27 = 729, the number of entries in the input data *N*. To improve the stability of our estimates, we lower the complexity of the model in two ways: limiting the range of error rates, and using parameter sharing.

**Limiting error rates.** We limit the error rates by requiring that 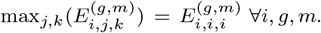. This guarantees that the largest fraction of the calls (*j*, *k*) are indeed correct (= *i*), separately for each error category.

**Parameter sharing.** We partition *t* into three disjunct subsets *t* = *t*_0_ ∪ *t*_1_ ∪ *t*_2_, where *t*_0_ = {(00, 00, 00)}, *t*_2_ = {(11, 11, 11)} and *t*_1_ = *t* \ (*t*_0_ ∪ *t*_2_), and require, whenever both *g*,*g*′ are in *t*_1_, that *θ*_*g*,*m*_ = *θ*_*g′*,*m*_ and 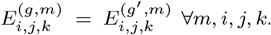. We choose this partitioning expecting that variants that are present in none (*t*_0_) or all (*t*_2_) of the genomes will be subject to different error rates than variants for which at least one family member is heterozygous (*t*_1_). These constraints allow us to introduce the notation *θ*_*s*,*m*_ := *θ*_*g*,*m*_ and 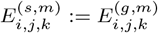 for all *g* ∈ *t*_*s*_ where *s* = 0, 1, 2, and reduce the number of algebraically independent parameters to 96. (See Supplementary A.1.2)

### 2.4 Inference

**Hidden components.** The product of Equations (2), (3) and (4) yields the joint distribution of observed (*G*^1^, *G*^2^) and hidden (*g*, *m*) variables,

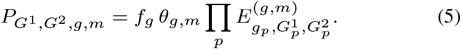

Under the assumption that the attributes (*G*^1^,*G*^2^,*g*,*m*) of different variants are independent and identically distributed (given *f*, *θ*, *E*), one can write the full and conditional distributions of the complete distribution 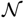 as (See Supplementary A.2)

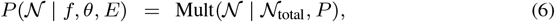

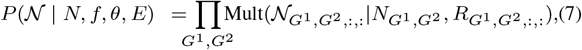

where 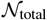 is the total number of variants (both in 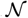 and *N*), 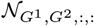 is a shorthand for 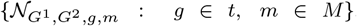, *P* = 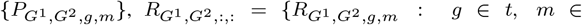 *M*} where 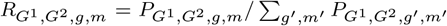 and 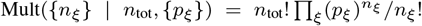 is the multinomial distribution. The distribution in Equation (6) describes an unconstrained sampling process where the variants can be freely distributed between bins with different (*G*^1^, *G*^2^,*g*, *m*) labels. The distribution of Equation (7) represents a constrained sampling process where the number of variants in each observable bin (*G*^1^, *G*^2^) are fixed to the actually observed value 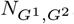, and only their hidden attributes (*g*, *m*) are subject to random sampling.

**Model parameters.** Using Bayes theorem, the conditional posterior of (*f*, *θ*, *E*) can be written as a product

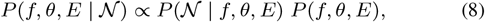

where the last term stands for the prior of the model parameters. Since 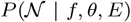 depends only on products of the model parameters (via Equation (5) and (6)), using a product of Dirichlet distributions as prior for (*f*, *θ*, *E*),
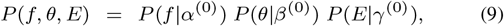

yields a posterior with identical structure (See Supplementary A.3),

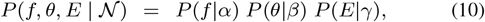

where, in Equations (9) and (10),

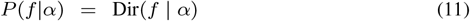

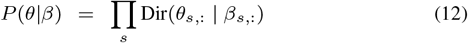

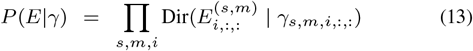

Here, Dir stands for the Dirichlet distribution, Dir(*x*|α) = 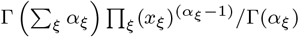 and its parameters are *α* = {α_*g*_ : *g* ∈ *t*}, *β* = {*β*_*s*,*m*_ : *s* ∈ {0, 1, 2}, *m* ∈ *M*}, γ = {γ_*s*,*m*,*i*,*j*,*k*_ : *s* ∈ {0, 1, 2}, *m* ∈ *M*, *i*, *j*, *k* ∈ *I*}. In Supplementary A.3, we show that the posterior values of *α*, *β, γ* are

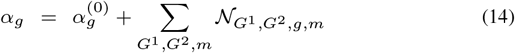

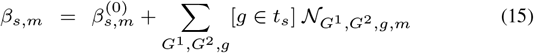

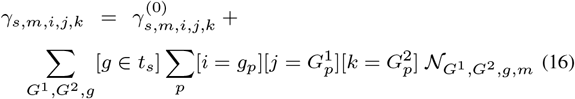

where [·] evaluates to 1 if the statement inside is true, and 0 otherwise.

**Gibbs sampling.** The conditional posterior of 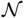 (see Equation (7)) and *f*, *θ*, *E* (see Equation (10)) are products of multinomial and Dirichlet distributions, respectively, both of which can be efficiently sampled. This enables Gibbs sampling, where we start with an initial guess (*f*,*θ*,*E*)^(τ=0)^. In step 1, we sample the independent multinomials of Equation (7) and obtain the τth sample of the hidden components, 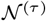. Then, in step 2, we sample the independent Dirichlet variables of Equation (10) and obtain the next sample of the model parameters, (*f*, *θ*, *E*)^(τ+1)^. We make sure that 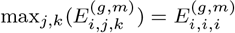 by first sampling each slice 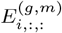 from an unconstrained Dirichlet distribution, followed by lowering all components that violate this criteria to the level of 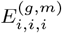 and re-normalizing. Repeating step 1 and 2 yields approximate samples from the joint posterior 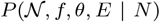. With appropriate “burn-in” (τ0) and “thinning” (Δτ), this procedure provides estimates of the hidden components in the form of a list of samples 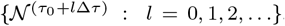. Details of choosing τ0 and Δτ are given in Supplementary B.

**Benchmarking metrics.** From the samples 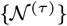, the τth estimate of any metric *μ* can be directly calculated, 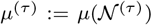, enabling accurate estimation of its posterior.

## 3 Materials and Methods

To assess the accuracy of our model, we carried out the validation experiment shown on Figure 3. First, we selected trios for which not only the NGS reads but also curated truth-sets of the children are published,and used two variant caller pipelines to produce variant calls from the read data. Second, we simultaneously used our model to estimate benchmarking metrics for both pipelines on all three members of the trio, and used the correct genotypes from the child’s truth set to calculate the true values of the same benchmarking metrics. Finally, we compared the estimated metrics with the true metrics to calculate the accuracy of our model.

**Fig. 3.**
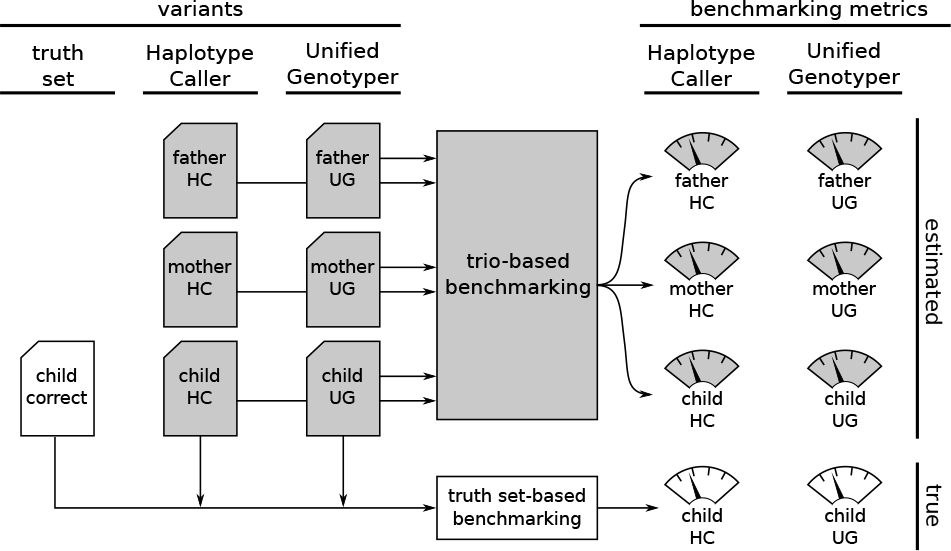
Validation experiment. We create two sets of trio variants called by a pipeline using GATK Haplotype Caller and another using GATK Unified Genotyper. Running our trio-based benchmarking model on this data produces estimates of various benchmarking metrics for both pipelines on all three data sets. At the same time, we use the correct genotypes for the child’s truth set to calculate the true values of the same benchmarking metrics. Gray shading marks the components that would be present even in a regular usage of our model, when the truth set is not available.

### 3.1 Data

Read data. We used 50x, 148-bp-long, paired-end, Illumina reads from three trios: the GIAB Ashkenazim trio^1^ (HG003, HG004, HG002), published by the Genome in a Bottle Consortium (Zook *et al.*, 2016), and 101-bp-long, paired-end, Illumina reads from the two unrelated trios from Platinum Genome^2^ (NA12889, NA12890, NA12877) and (NA12891, NA12892, NA12878), published by Illumina, Inc. (Eberle *et al.*, 2017). From these alignment files, we extracted and sorted the reads using samtools^3^, and dropped duplicates and unpaired reads, using a custom script. Command lines are listed in Supplementary D.

**Truth set for child.** Curated, high-confidence variants are publicly available for all three children, HG002^4^, NA12877 and NA12878^5^. We used bcftools^6^ to select bi-allelic SNPs on chromosomes 1-22 (~ 4 million). Merging and comparing with the variants produced by the variant caller pipelines allows direct, “truth set-based”, benchmarking of the pipelines. (See Supplementary D for command lines.) We restricted the scope of our validation experiment to SNPs to avoid having to reconcile different representations of identical variants, which often arise for indels.

**Pipelines.** We used two variant caller pipelines, both of which uses BWA-MEM 0.7.13 (Li and Durbin, 2009) for alignment, GATK tools (McKenna *et al.*, 2010) for local realignement and base quality recalibration, and take advantage of known SNPs and indels (HapMap, dbSNP, 1000GP, Omni) according to GATK Best Practices recommendations (DePristo *et al.*, 2011; Van der Auwera *et al.*, 2013). In the variant calling step, one pipeline used GATK HaplotypeCaller v3.5, and the other GATK UnifiedGenotyper v2.7. Command lines are listed in Supplementary D.

**Aggregating genotype calls.** For each trio, we merged the SNPs called by the two pipelines using bcftools, and filtered out the variants that fell outside the high-confidence regions or become multi-allelic. We aggregated the remaining SNPs (~ 4 million) by counting the occurrences of each (*G*^1^, *G*^2^) combinations, where the trio genotype *G*^1^ is called by Haplotype Caller, and *G*^2^ by Unified Genotyper. During aggregation, we recorded missing variants as genotype 00 and disregarded phasing information, i.e. regarded both heterozygous genotypes, 0|1 and 1|0, as 01. This yielded the observed counts of joint trio calls 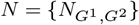.

**Subsampling.** To test the robustness of our model, we repeated the validation experiments for subsets of SNPs of different sizes. We subsampled a pre-defined number of variants from *N* by choosing a random subset of SNPs with uniform probability.

### 3.2 Running the estimator

**Imputing “uncalled variants”.** Neither Haplotype Caller nor Unified Genotyper reports homozygous reference genotypes (i.e. 00) when run on a single sample. Due to this limitation, the number of potential variants that do not get called 01 or 11 by either pipeline in any family member, i.e. 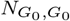 where *G*_0_ = (00, 00, 00), is unknown. Although the maximum number of loci available for SNPs is the total length of the genome, not all imaginable SNPs are equally plausible, and most likely not all loci are subject to the same genotyping error model. Instead, we impute 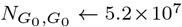, the total number of variants reported in the dbSNP database^7^, which is a plausible maximum number of potential variants among all human genomes.

**Details of Gibbs sampling.** For each set of input data N, we perform Gibbs sampling as described in Section 2.4. We set the parameters of the Dirichlet priors to 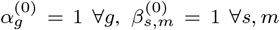 and 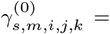 = 1000, if *i* = *j* = *k* and 1 otherwise ∀_*s*,*m*,*i*_. The skewed choice for γ^(0)^ represents our expectation that the error rates are small 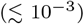. We draw the initial values of *f*, *θ*, *E* from their prior, and let the Gibbs sampler run for τ_0_ = 50,000 iterations, allowing it to reach equilibrium. We proceeded by running 100,000 iterations, while recording samples with Δτ = 1000 period, to decrease the correlation between consecutive samples. This procedure resulted in 100 samples, 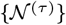, drawn from its posterior 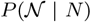. With our scipy-based Python implementation, this procedure takes 4.5 hours, using 1 CPU on the AWS instance m1.small.

### 3.3 Metrics

For each trio, we run the trio-based benchmarking, which produces samples 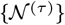. From this, we directly calculate the samples of the confusion matrix for the child {*n*^(child)(τ)^} (from Equation (1)). Using the formulas for precision and recall, we obtain their samples {prec^(τ)^} and {rec^(τ)^}, for both pipelines. We estimate each metric with the average of its samples.

To investigate how sensitive our model is to the total number of variants, we repeatedly sub-sample the observed counts *N*, and run the trio-based benchmarking all over again. We do this for 8 different number of variants (10^3^ – 3 × 10^6^), repeating each sub-sampling and estimation 10 times for every trio.

## 4 Results

The simplest method of analyzing trio results from the two pipelines is counting Mendelian violations. This, alongside with the true genotyping errors in the child, is shown in Table 1. Comparing the two columns shows that the difference in the number of Mendelian violations correctly indicate the difference in the actual number of genotyping error between the two pipelines.

**Table 1.** Number of genotyping errors in the children’s genomes, calculated from comparisons with the truth sets, and the number of Mendelian violations in their respective trios. Counts are shown for Haplotype Caller and Unified Genotyper.

|  | Genotyping errors in child |  | Mendelian violations in trio |  |
| --- | --- | --- | --- | --- |
|  | HC | UG | HC | UG |
| HG002 | 7,496 | 20,142 | 3,946 | 7,601 |
| NA12877 | 36,759 | 39,780 | 57,179 | 59,038 |
| NA12878 | 34,436 | 37,224 | 11,875 | 14,286 |

To see how accurately our trio-based benchmarking method can compare the two pipelines, we compare estimates of precision and recall with their true values for the children in Table 2. Although the estimated values of the metrics significantly differ from the truth, both the signs and magnitudes of *differential* metrics are accurately estimated. This is also visible from the smaller estimation uncertainty σ_trio_, reported by the model.

**Table 2.**
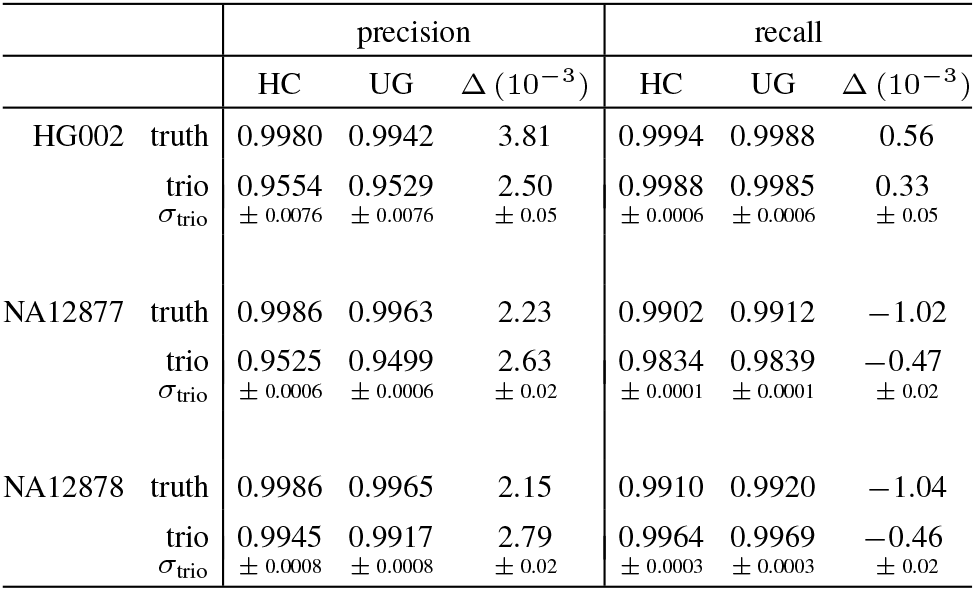
Precision and recall of the two pipeline (HC: Haplotype Caller, UG: Unified Genotyper) on the children’s samples calculated by truth set-based benchmarking, and estimated with trio-based benchmarking, using all 4 million SNPs. Δ denotes the difference between HC and UG values. otno is self-reported uncertainty of the model, i.e. standard deviation of the Gibbs samples.

In Figure 4, we plot the deviation of the trio-based estimate from the true value as a function of number of variants. While the estimation error of the metrics themselves hovers around 0.1-0.01 even when large number of variants are considered, the model’s error on the differential metrics is around 10^−3^ for large number of variants, and stay below 5 × 10^−3^ even for a mere 1000 variants. Recall values (both absolute and differential) are estimated more accurately then precision.

**Fig. 4.**
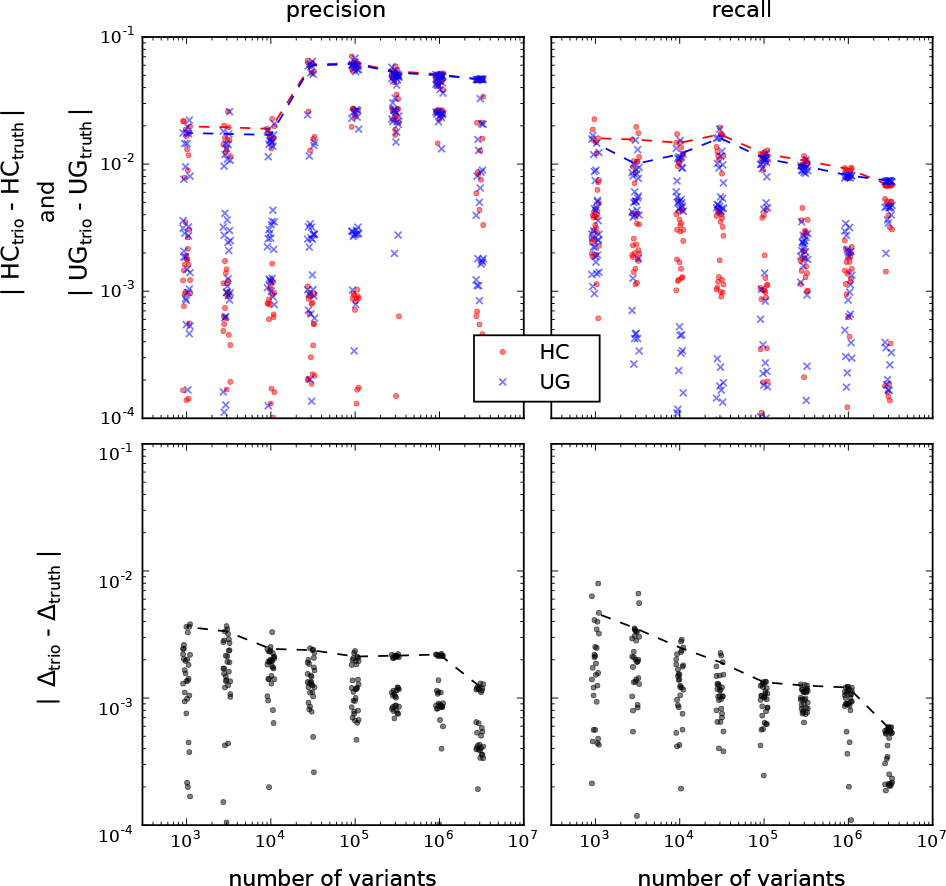
Estimation errors. **Top:** Difference between estimated (from trio) and true values (from truth set) of precision and recall. **Bottom:** Difference of the estimated and true differential precision and recall. Markers show the results obtained from the validation experiments run on sub-sampled variants. Dashed lines highlight the 90th percentile of each group.

To understand the reason why our model is much better at estimating differential metrics than the metrics themselves, we plot the entries of the genotype confusion matrix (*n*) for NA12878 in Figure 5. Our model accurately estimates the number of loci where either one of the pipelines makes an incorrect call, but not both. Since these entries of the confusion matrix contribute the most to differential performance, it is no surprise that the *difference* in precision and recall are accurately estimated, even if their actual values are not. We can intuitively understand this disparity in the following way: We expect the two pipelines to have similar performance. Only the 27 diagonal elements (*G*^1^ = *G*^2^) of *N* convey information about this common performance. On the other hand, all off-diagonal elements (*G*^1^ ≠ *G*^2^) tell us something valuable about the differential performance. Since there are more of the latter, they paint a more detailed picture, and allow us to estimate the differences more accurately.

**Fig. 5.**
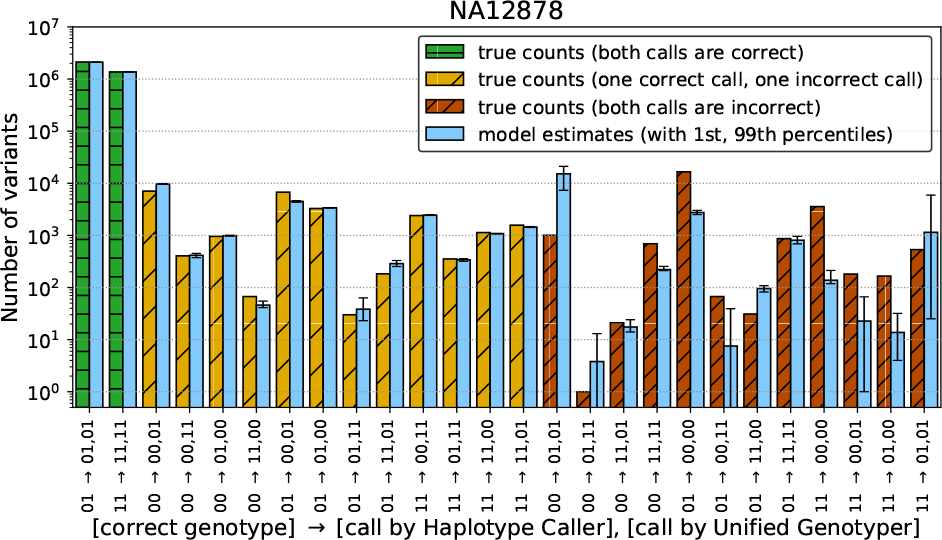
Genotype confusion matrix of NA12878. True values are shown alongside with the estimates obtained from trio-based benchmarking. Error bars on the estimates indicate 1st and 99th percentiles of the samples from the Gibbs sampler. (See Supplementary Figure S.6 for HG002 and NA12877.)

## 5 Discussion

Aggregating observed Mendelian violations across many loci in related individuals provides an approximate notion of the accuracy of variant calling pipelines. When comparing two pipelines, we expect the better one to produce fewer Mendelian inconsistencies, and indeed, as shown in Table 1, this is the case: Haplotype Caller produces fewer Mendelian violations in all three trios, and makes fewer genotyping errors for all three children.

In the light of our results in Table 2, two caveats of this naive method are immediately visible: First, the numbers of Mendelian violations do not accurately reflect *how much* better is the better pipeline. Second, the bare counts of Mendelian violations do not even hint that Unified Genotyper actually has a higher recall on the samples NA12877 and NA12878, something our trio-based benchmarking method estimates correctly (See Table 2). This exemplifies that estimating precise quantitative differences between variant caller pipelines, in the absence of a reliable truth set, is a non-trivial task. As we demonstrated, our method can do this with high accuracy.

Previous trio-based benchmarking approaches range from fitting a calibration curve between observed number of Mendelian violations and true number of errors (Hao *et al.*, 2004), to modeling haplotypes and recombination events during meiosis (Markus *et al*., 2011; Kojima *et al*., 2013; Peng *et al.*, 2013). All previous approaches focus on a set of variants produced by a single pipeline, disregarding all information about the correlation between pipelines. We showed that by analyzing the joint counts (*N*) of the called genotype trios from two pipelines one can estimate differential performance with high accuracy (with uncertainty of 10^−3^). In fact, exactly because of the information contained in the correlations, the differential metrics can be estimated with more than an order of magnitude lower uncertainty than the metrics themselves. As a consequence, our uncertainty is an order of magnitude lower than it was previously achieved with dyads (Haaland and Skaug, 2013), and comparable to the uncertainties obtained from pure simulations for trios (Hao *et al*., 2004).

Although our current method is limited to using bi-allelic variants from trios, the mathematical framework enables straightforward generalization. Data from multi-allelic variants and larger pedigrees can be incorporated by expanding the sets *I* and *T*. The demand of computational resources, which increases exponentially with the sizes of *I* and *T*, can be curbed by truncating the model to limit the number of genotyping errors per locus, which enables implementation with sparse arrays.

The estimated complete distribution 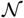 and the person-specific confusion matrices *n*^(*p*)^ contain a wealth of additional information that can be directly used to *improve* the performance of variant calling. Bayesian model averaging (See e.g. Fragoso and Neto (2015)) can be used to reconcile genotypes at loci where the pipelines do not agree. By counting how often one pipeline is right when the other is wrong for each called trio combination (*G*^1^, *G*^2^), one can estimate the posterior probability of the correct genotype, *P*(*g* | *G*^1^, *G*^2^). E.g. from the confusion matrix *n* shown on Figure 5, we can calculate the probability of HC being correct when HC and UG call 01 and 00, respectively: *P*(01 | 01, 00) = *n*_01, 01, 00_/∑_*i*∈*I*_ *n*_*i*,01,00_ (≈ 0.72). One can imagine extending this scheme to a council of pipelines where, instead of giving equal say to each pipeline, a weighted voting is carried out to determine the genotype of each variant.

1 ftp://ftp-trace.ncbi.nlm.nih.gov/giab/ftp/data/AshkenazimTrio/

2 ftp://ftp.sra.ebi.ac.uk/vol1/ERA172/ERA172924/bam/

3 https://github.com/samtools/samtools

4 ftp://ftp-trace.ncbi.nlm.nih.gov/giab/ftp/release/AshkenazimTrio/(v3.3.2)

5 ftp://platgene_ro@ussd-ftp.illumina.com/2016-1.0/hg19/small_variants/

6 http://samtools.github.io/bcftools/

7 https://www.ncbi.nlm.nih.gov/projects/SNP/

## Acknowledgements

Discussions with Yilong Li, Lizao Li, Maxime Huvet, Amit Jain, Milos Popovic, Aqeel Ahmed, Maria Suciu, Sun-Gou Ji, Ivan Johnson, Kaushik Ghose, İlker Gündoğdu were very valuable at multiple stages of this work. We are grateful to them for their insights and suggestions.

